# Mountable miniature microphones to identify and assign mouse ultrasonic vocalizations

**DOI:** 10.1101/2024.02.05.579003

**Authors:** Elena N. Waidmann, Victor H.Y. Yang, William C. Doyle, Erich D. Jarvis

## Abstract

Vocal communication is a major component of animal social behavior. Vocalizations can be learned or innate, and can convey a variety of signals, including territorial limits, the presence of predators, or courtship intent. Mouse ultrasonic vocalizations (USVs) are a promising model in which to study mammalian vocal production circuits. While mouse USVs are innate, mice still show complex vocal behavior, including production of structured song composed of multiple syllable types and the ability to modify their vocal rate and syllable repertoire based on social conditions. Though in courtship interactions male mice produce the majority of the emitted USVs, female mice are capable of emitting USVs. In order to study the underlying mechanisms of vocal production in freely behaving pairs of mice, it is necessary to identify the individual responsible for each syllable in group settings. Previous methods to identify the source of an individual USVs have used high-density microphone arrays and triangulation methods, which involve the use of multiple costly microphones and require implementation of complex computational methods. Here we identified, developed, and used an inexpensive, mountable, ultrasound-sensitive miniature-microphone system to record and identify USVs from individual mice during dual socializing behavior. Our system includes custom circuit boards that can be fitted to individual mice and connected to a variety of existing USV recording systems. We found that these miniature microphones reliably detected mouse USVs, and that a high percentage (90%) of vocalizations could be attributed to a specific animal in a vocalizing pair based on the relative amplitude differences alone. This simple readout method avoids the implementation of complicated triangulation methods. By pairing this method with simultaneous video recording and automated animal body part and identity tracking, we were able to study and describe the broader courtship behavioral landscape, in which USV production is one component. These results offer a promising, low-cost, and simple method that researchers can implement to study the social vocal communication between at least pairs of vocalizing mice.

## Introduction

Vocalization is a widespread form of communication across the animal kingdom, used to convey information about predators, reproductive fitness, and courtship readiness. The house mouse, mus musculus, produces both audible squeaks in response to aversive stimuli and social ultrasonic vocalizations (USVs) to communicate with conspecifics [1]. USVs are composed of discrete syllables, formed into longer ‘song’ bouts, and are mostly innate [2,3], produced from birth by pups to initiate maternal retrieval when separated from the nest [4,5]. After a period of juvenile development in which mice rarely produce USVs, young adults begin to produce social USVs [6]. In particular, male mice readily vocalize for long bouts during female-directed courtship [1], and these vocalizations often induce females to approach the males [7,8]. Males do demonstrate some flexibility in their vocal usage and repertoire depending on social context [8], and signatures of individual identity across different individuals [9], which suggests that USV production is a dynamic and complex social behavior. While much of the study of mouse ultrasonic vocal behavior is centered on male courtship song, females are also capable of producing USVs, both in male-female [10,11,12] and female-female interactions [13].

Prior research in USV production behavior has focused mostly on male vocalizations, and may have underestimated the role that females play in these social interactions, both in their production of USVs, and in their behavioral response to external USVs. In order to study the complex behavioral interactions between two or more vocalizing animals, it is necessary to parse which vocal syllables are produced by which individual, which is impossible with a traditional recording setup, using a single overhead microphone (Fig 1a). USVs are produced at close range [11] and at high frequency [14]. Recently, to overcome this credit-assignment problem, several labs have developed array systems of microphones surrounding or above a recording arena [10,11,12]. Complex mathematical and computational tools are then used to sort out which vocalizations belong to whom. While these methods increase animal identification fidelity, it is expensive to multiply high-fidelity microphones in a single recording arena, as well as difficult to computationally and analytically triangulate the source of individual USVs, especially those produced when animals are in close proximity. In other vocalizing species, particularly songbirds [15] and bats [16], researchers have developed wearable microphones that individuals carry and whose amplitude differences can be used to assign the identity of a single vocalization. However, these existing technologies are designed to detect low-frequency vocalizations, and do not capture mouse vocalizations in the ultrasonic range (30 - 100 kHz). We sought to develop similar wearable microphones for mice, in order to simplify the credit-assignment problem for mouse USVs.

**Fig 1.**
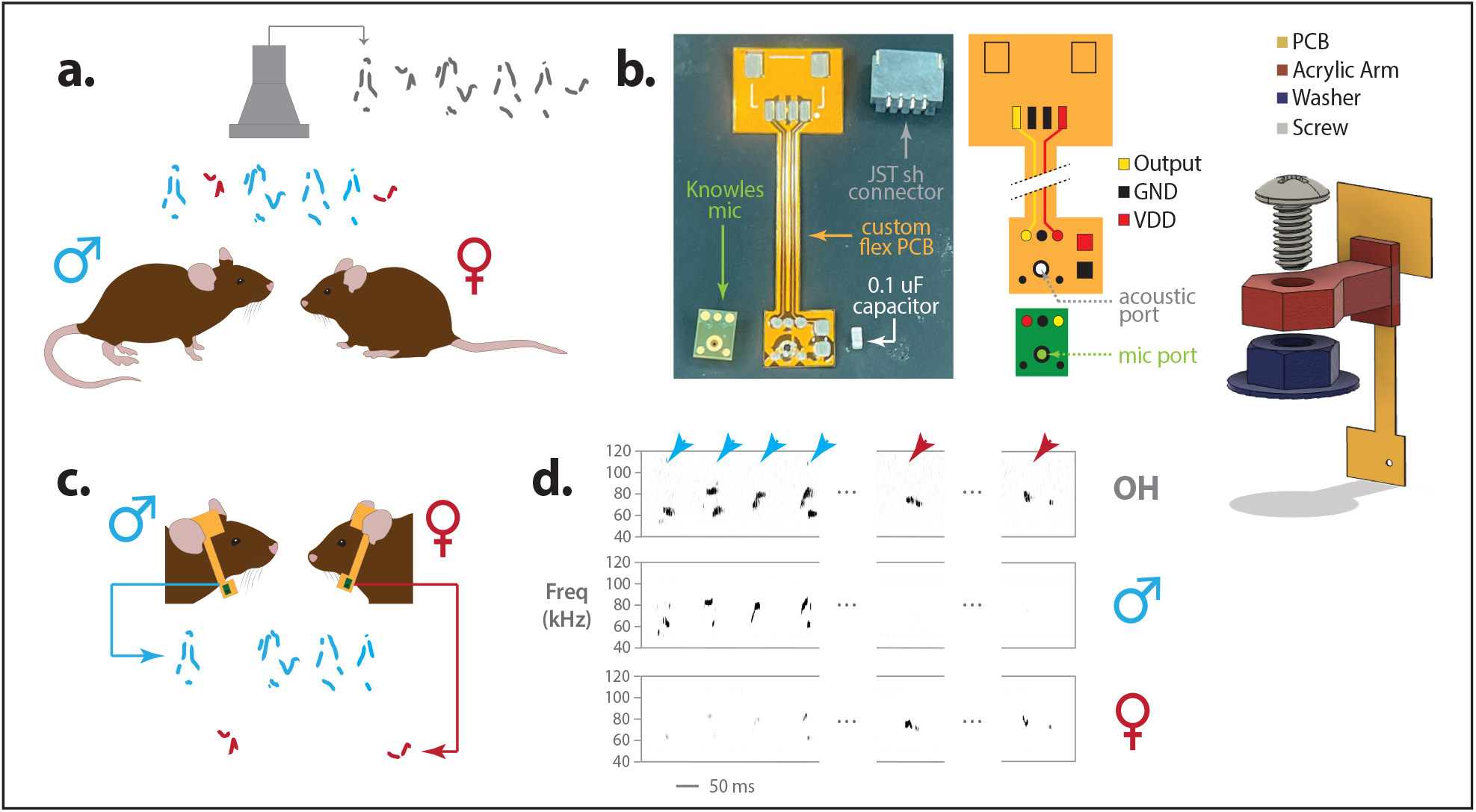
Wearable mini-microphone design and recording in social pairs. (a) Identity assignment problem during multi-animal mouse ultrasonic vocalization (USV) sessions. A single ultrasonic overhead microphone used to record USVs in a social context (gray) cannot discriminate between syllables that originate from the male (blue) or the female (red). (b) Design and layout of the mini-microphone recording device. Left, photograph of the mini-microphone circuit board and JST connector against a blue paper background. Right, 3D model of the custom attachment arm, consisting of the acrylic arm (red) secured between a washer (blue) on the mouse’s skull and a screw (gray). (c) Schematic of male and female mice with attached mini-microphones, each able to parse out vocalizations from the individual mouse. The printed circuit board (yellow) is glued to the acrylic arm, with the mini-microphone acoustic port pointed towards the mouse’s mouth. (d) Segments of spectrograms from mini-microphone recordings, showing vocalization assignment. Top, recording from overhead (OH) microphone collecting all vocalizations. Middle, recording from mini-microphone attached to the male mouse. Bottom, recording from mini-microphone attached to the female mouse. Amplitude differences between male and female spectrograms were used to assign individual vocalizations as male (blue arrows) or female (red arrows).

Here, we identified an ultrasound-capable miniature microphone, and developed custom hardware and credit-assignment analysis. All together, the system is very low-cost and easy to implement. It is also customizable and can integrate into existing recording systems. We used this system to record extended courtship sessions in multiple pairs of freely behaving mice, and demonstrate that our system can, with simple visual and computation analysis, assign a high percentage, 90%, of USVs to individual animals. We confirm that females do produce USVs readily and throughout extended social interactions with males, but at quite low rates. Finally, we integrate the vocal behavior with video data to identify differences in animal positions during the production of male and female vocal interactions to build a larger picture of mouse social behavior.

## Results

We attempted to solve the vocalization credit-assignment problem in a high-fidelity manner, while minimizing cost and computational investment. We identified a commercially available small (2.75mm length x 1.85mm width x 1.00mm height) and lightweight (0.03 g) micro-electrical mechanical system (MEMS) microphone, normally used for detecting high-frequency sounds to diagnose problems in electronics or buildings (Knowles SPU0410LR5H-QB; Fig 1b).

We first found that the mini-microphone was capable of detecting mouse USVs played from an ultrasonic speaker (Avisoft BioAcoustics, Ultrasonic Dynamic Speaker with 216H US Playback interface), played at the same decibel (loudness) level that mice normally produce USVs (Fig 2a). Importantly, when the angle of the mini-microphone was changed from directly parallel to the speaker to 30° to 180° degrees to the speaker, the playback vocalizations were much less detectable (Fig 2b,c,d). The detection level dissipated at a distance of about 10 cm from the speaker (Fig 2e), within the normal distance ranges in mice in paired conditions. Distance detection decreased with decreasing amplitude (Fig 2f,g). These findings suggest that the ultrasonic mini-microphones are very sensitive to the angle and distance from which the sounds originate.

**Fig 2.**
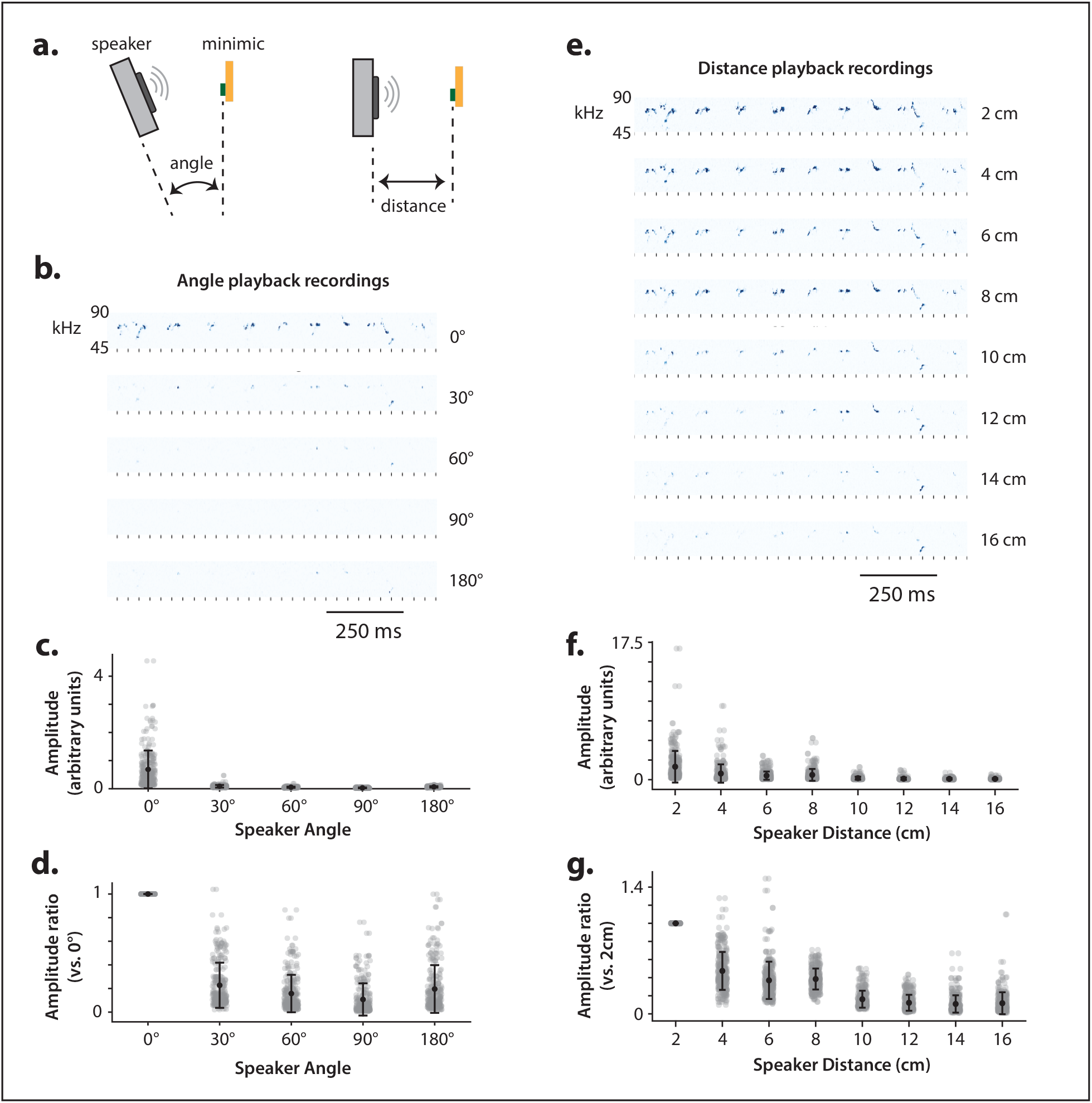
Distance and angle sensitivities of the miniature microphones. (a) Schematic of the mini-microphone distance and angle tests. An audio file of recorded USVs was played on an ultrasonic speaker (gray) and recorded on the mini-microphone (yellow) at varying angles and distances between the speaker and microphone. (b) Spectrogram segments taken at different recording angles (0° to 180° at 30° increments) at a distance of 6cm. A 1-second excerpt for the recording is shown. Amplitude visibly drops off with greater angle deviation. (c) Amplitude distribution at each recorded angle. Each dot represents the amplitude of a single syllable. Black line, mean ± 1 standard deviation of amplitude distributions for each angle. (d) Mean amplitude ratios at each angle relative to the 0° angle. For each individual syllable, at each angle, syllable amplitude was averaged within the time range identified on the 0° recording. For every syllable, the amplitude at angle *a* is divided by the amplitude of that same syllable at the 0° angle (ratios are all 1 at 0°). Black line, mean ± 1 standard deviation. (e) Spectrograms at different recording distances, between 2 cm and 16 cm at 2 cm increments. The 1-second excerpt and spectrogram range are the same as those shown in Fig 2b. Amplitude visibly drops off with distance. (f) All syllable amplitudes at each recording distance; data is shown in the same format as that of Fig 2c, with angle parameters replaced by distance. (g) Comparison of each syllable amplitude between 2cm and each other distance. Analysis is conducted with the same procedure as angle in Fig 1d but relative to the 2cm distance.

We next attached the mini-microphone to a custom printed circuit board (PCB), which in turn we attached to a 3-D printed mounted headgear; the audio signal was transferred through a micro-JST connector to an Ultrasound Gate 416H (Avisoft) recording system (Fig 1b). The total weight of the microphone, circuit, and headgear mount is 1.17g, over 63g lighter than the most commonly used overhead ultrasonic microphones (Avisoft Bioacoustics CM24/CMPA condenser microphone). We then mounted the mic headgear to a screw mount on the mouse head (placed earlier during surgery) for each individual mouse with the acoustic port close to 0° to the mouth (Fig 1c) and thus 90-180° to another animal if face-to-face.

We recorded 9 naturalistic, free behavior courtship sessions, each between one male and one female mouse, each with a mounted mini-microphone, ranging from 20 to 40 minutes in duration. We used 4 males and 2 females, with 3 pairs recorded twice. In this set up, we were able to record male and female USVs, and distinguish amplitude differences between the mics (Fig 1d). We collected 22,561 USV syllables in total (full list of individual pairings and USVs per pairing is in Table 1).

**Table 1.** Distribution of USV assignment in male and female pairs using the mini-microphones

| Session (M_F) | Total # USVs | # Male USVs | # Female USVs | # Unassigned USVs |
| --- | --- | --- | --- | --- |
| B770_B744 (1) | 6691 | 5821 (87.0%) | 178 (2.7%) | 692 (10.3%) |
| B757_B744 | 621 | 553 (89.0%) | 32 (5.2%) | 36 (5.8%) |
| B770_B745 | 3688 | 3283 (89.0%) | 77 (2.1%) | 328 (8.9%) |
| B782_B744 (1) | 2575 | 2420 (94.0%) | 26 (1.0%) | 129 (5.0%) |
| B782_B744 (2) | 1248 | 690 (55.3%) | 303 (24.3%) | 255 (20.4%) |
| B782_B745 (1) | 1983 | 1593 (80.3%) | 135 (6.8%) | 255 (12.9%) |
| B770_B744 (2) | 857 | 551 (64.3%) | 40 (4.7%) | 266 (31.0%) |
| B782_B745 (2) | 2740 | 2487 (90.8%) | 76 (2.8%) | 177 (6.5%) |
| B805_B745 | 2158 | 1443 (66.9%) | 389 (18.0%) | 326 (15.1%) |
| **TOTAL** | **22561** | **18841 (83.6%)** | **1256 (5.6%)** | **2464 (10.9%)** |

We also recorded concurrent video data, in order to align the recorded vocal behavior with broader social and courtship behavioral motifs. To perform these video recordings, we designed and constructed a recording box from opaque white acrylic, with overhead openings for an ultrasonic microphone, overhead camera, and mini-microphone cables (Fig 3a). The box had a clear acrylic base, and was suspended on an external aluminum frame, to allow a recording from an additional bottom-view camera (Materials and Methods). We used a manual cable rotator to avoid tangling between the mini-microphone cables of the two freely-moving animals.

**Fig 3.**
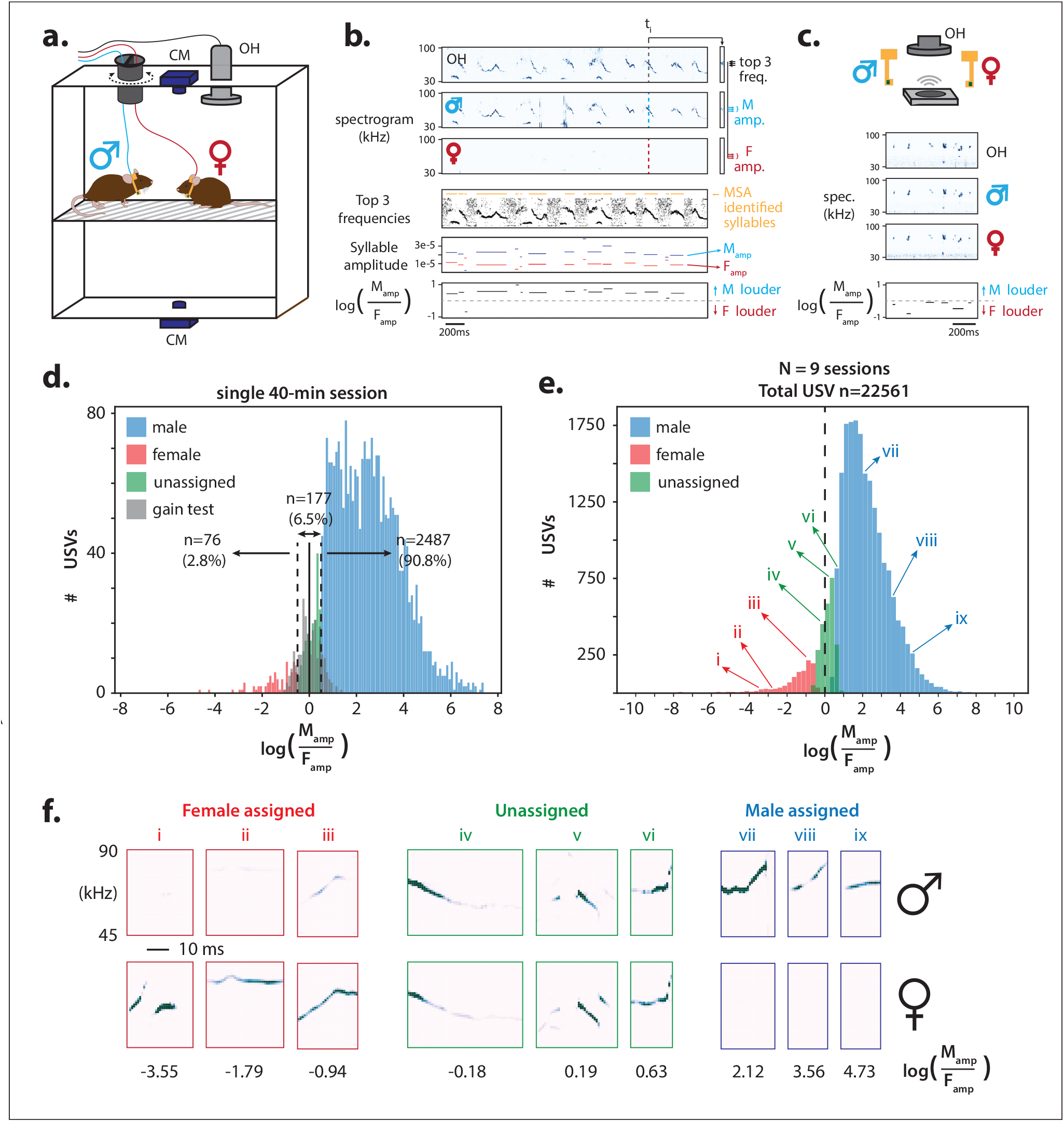
Behavioral set up and vocalization assignments. (a) Schematic of custom behavioral rig for paired social recordings (male ♂, blue; female ♀, red). Each mouse wears a mini-microphone (yellow), and are allowed to freely interact in an acrylic box (white) on a clear acrylic floor (gray dashes). An overhead ultrasonic microphone (gray, “OH”) records all vocalizations that occur, while top and bottom cameras (blue, “CM”) record movement. Mini-microphone cables (red, blue) are run through a manual cable untangler (black). (b) Computing male-female amplitude differences. Top 3 rows: spectrograms of recordings from the overhead, male mini-microphone, and female mini-microphone respectively. 4th row: The top 3 frequencies from the overhead recording, with start and end times of each syllable identified using MSA 2.0 (orange). 5th row: Median male mini-microphone amplitude (blue) and female mini-microphone amplitude (red), averaged across the time and frequency segments shown in row 4. 6th row: log of male:female amplitude ratio between two individuals. (c) Pre-session gain-calibration. Top, schematic of speaker playing USVs to overhead, male, and female microphones. Spectrograms demonstrate signals of roughly equivalent amplitude across all three recording channels. As in Fig 3b, a male mic:female mic amplitude ratio is calculated for each syllable. (d) Histogram displaying the male:female amplitude ratios of all syllables across a single 40-minute recording session. The distribution of the ratios from the gain-matching session (gray, solid red line: mean; ± 1 standard deviation: dashed red line) was used as the point of comparison for amplitude differences in the courtship session. Male-assigned (blue), female-assigned (red), and unassigned syllables (green) based on difference from gain-matching session mean. (e) Histogram showing male:female amplitude ratios zeroed to the respective gain-matched mean ratio, for 22561 USVs across 9 sessions. The majority of syllables are louder on the male mini-microphone (rightward skew to the red line). (f) Examples of sonograms from female-assigned (red), unassigned (green), and male-assigned USV syllables and amplitude differences between mini-microphones of a male and female pair. Top row, recording from male mini-microphone; bottom row, recording from female mini-microphone. Black numbers show male:female amplitude ratios, roman numerals show position on amplitude ratio distribution in Fig 3e.

With the overhead microphone and respective mini-microphones, a total of 3 sound channels were recorded during each session and converted to spectrograms (Male, Female, and Overhead). We used the overhead microphone as a ground-truth record for all emitted vocalizations. To compute syllable amplitude, for each spectrogram timepoint, we identified the 3 frequencies with the highest amplitude on the overhead channel to avoid capturing any mechanical or electrical noise that may be present. We computed the mean amplitude on the male and female channels across those frequencies for each timepoint (Fig 3b). We performed syllable identification and identified the start and end of each USV syllable on the recording from the overhead microphone using the Mouse Song Analyzer 2 (MSA2) function of Analysis of Mouse Vocal Communication (AMVOC) software [17]. By measuring the median amplitude on the separate male and female recording channels across the duration of syllables, we obtained a median male syllable amplitude (*M*_amp_) and median female syllable amplitude (*F*_amp_) for each USV. Finally, we took the log of the ratio between male and female syllable amplitudes,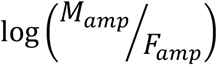, to generate an amplitude-ratio metric for each syllable (Materials and Methods).

Because the sensitivity of individual mini-microphones may vary across sessions, the amplitude ratio alone is not sufficient to assign a single USV as either male-or female-emitted. To compensate for these inherent variations, we ran a brief microphone calibration procedure with USVs played from the ultrasonic speaker, equidistant to each different microphone, prior to each recording (Fig 3c). Gains of both mini-microphones were manually adjusted such that a playback recording of mouse USVs had similar amplitude. We recorded 30-60 seconds of the playback on all three recording channels before each male-female recording session. We then computed the male-female amplitude ratio (as above, Fig 3c) for each syllable recorded in the gain test session. We used the distribution of these gain-matched amplitude ratios as a baseline against which to measure male and female vocalizations produced during the social recording. If a syllable in the social recording had a log 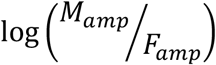 greater than 1 standard deviation above the mean amplitude-ratio from the gain test, we assigned it as male; if log 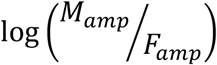 was less than 1 standard deviation below the gain test mean, we assigned it as female; and if it fell within 1 standard deviation of the gain test mean, we designated it as unassigned.

An example of this process shows the distribution of amplitude ratios for a single 40-minute male-female session, in which 90.8% of vocalizations were assigned as male-emitted, 2.8% as female-emitted, and the remaining 6.5% were unassigned (Fig 3d). Across all 9 male-female sessions of 22,561 unique vocalizations, 83.6% (range 55.3-94.0) were assigned as male-emitted, 5.6% (range 1.0-24.3) as female-emitted, and 10.9% (range 5.8-20.4) were unassigned (Fig 3e, Fig S1, Table 1). Fig 3f shows spectrograms of example male, female, and unassigned syllables from across the range of amplitude ratios. Thus, this analysis demonstrates that although in novel male-female pairings, the vast majority of USVs are produced by the male, females produce a range of USVs as well.

We then sought to integrate our audio recordings and credit assignment methods with the video data, to test the utility of our system for studying holistic vocal and courtship behavior in freely behaving animals. For 6 of the male-female courtship sessions, we recorded and aligned video of the mice with the audio output. We trained a SLEAP animal pose tracking model [19] to track various body parts on each mouse (Fig 4a). We computed the distance between the body center point of each animal as ‘inter-animal distance’. We found that USVs were produced primarily when animals were in extremely close proximity (most often 0 to 10 cm apart, Fig S3a), and occurred in repeated bouts throughout the length of the 40-minute sessions (Fig 4b, top). By using our credit-assignment system, we identified the individual producing each vocalization, and could therefore calculate the inter-animal distance during male, female, or unassigned USVs. We found that while both males and females most often produced USVs when they were very close to each other (less than 10 cm), there were sex differences where males produced USVs within the entire distance range (some even at nearly 40 cm apart), but that females mainly produced USVs when the males were very close to them (example in Fig 4b, bottom). These distance profiles and relationships were significant, where across all 6 pairs, the mice were significantly closer during USV versus non-USV production periods (ind. t-test; voc. vs nonvoc. (rand) p=0, voc. vs nonvoc. (all) p=0, nonvoc. (rand) vs nonvoc. (all) p=0.731)), and female and unassigned vocalizations were produced at significantly closer distances than male vocalizations (ind. t-test; M vs F p=1.08E-07; M vs F p= 3.11E-10; M vs UN p=0.403; Fig 4c, Fig S2). To better define body positions during USV production behavior, we also measured the distance from the male nose to the female nose and to the female tail base during only ‘close’ USVs (<8cm, about the length of an adult mouse), which accounted for 68.2% of all USVs, (Fig S3a). We found that male vocalizations most commonly occurred when their nose was in extremely close proximity to the female ano-genital region, while female and unassigned vocalizations were more frequently produced when the animals were nose-to-nose (Fig 4d). These findings suggest that males are more motivated to produce courtship USVs when chasing females with ano-genital olfaction or stimulation, whereas females are motivated to produce USVs face to face with males.

**Fig 4.**
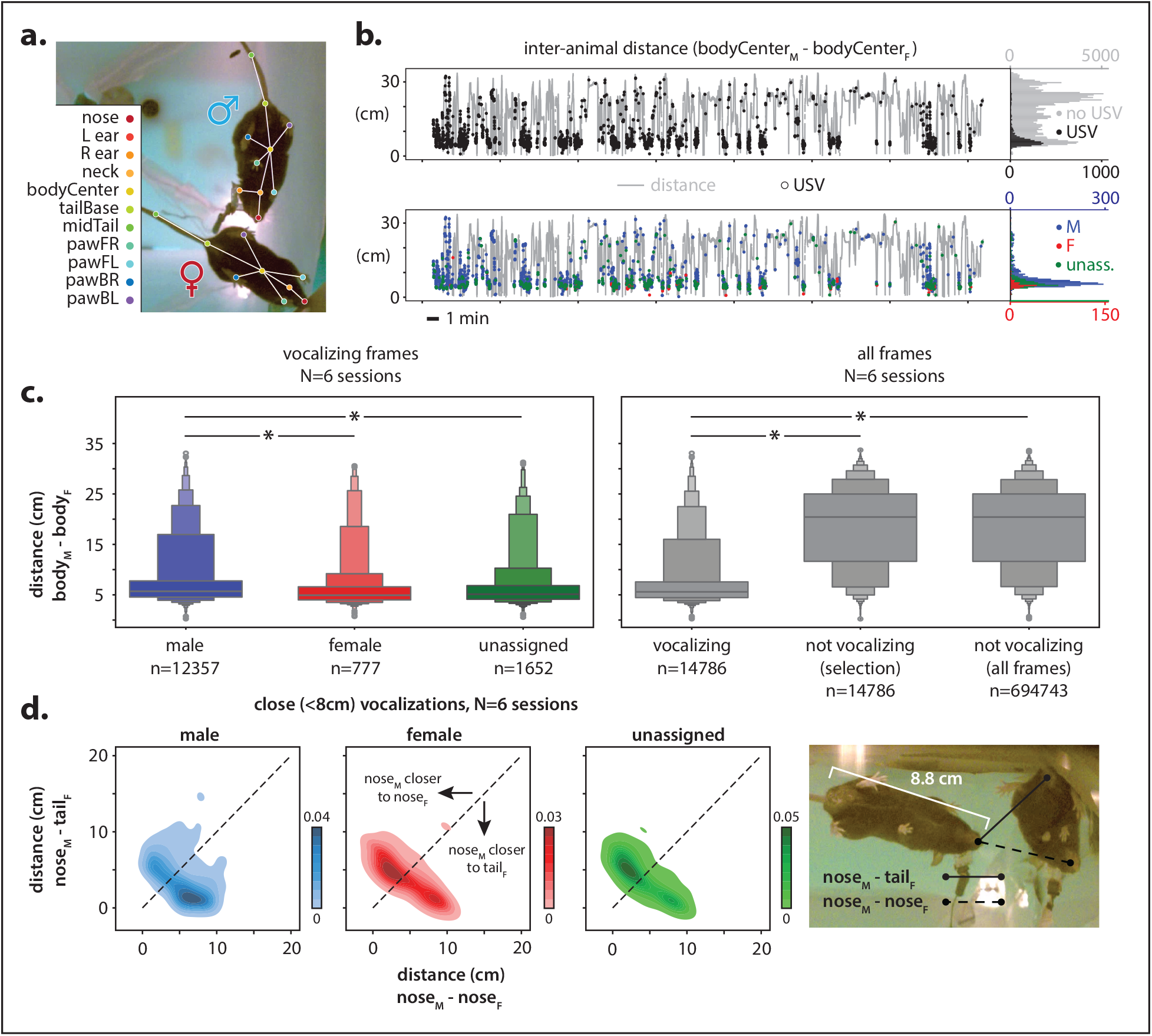
Integrating vocalization and movement with video tracking. (a) Tracking mouse body parts during video recording using SLEAP. A total of 11 separate body features were defined and manually labeled. (b) Inter-animal distance across a single recording, measured male bodyCenterM to female bodyCenterF. Top: video frames in which any USV was detected are labeled with black dots. Bottom: video frames in which male (blue), female (red), or unassigned (green) syllables are labeled with colored dots. Right of each distance plot, histogram of distribution of distances during occurrence of each type of vocal behavior. (c) Comparing inter-animal distance across vocalization types. Left: plots showing distribution of body distance between animals during frames containing male, female, or unassigned vocalizations. Right, plot of all video frames showing distributions of distance for all vocalizations, an equivalent number of randomly-selected non-vocalizing periods, and all non-vocalizing periods. Data shown aggregates six 20-40-minute recordings with both audio and video data. (d) Left: Density plots comparing noseM-noseF and noseM-tailF distance for each close (≤ 8 cm) USV. Unity line, dashed black line. Right: example of noseM-noseF and noseM-tailF distances.

## Discussion

In this study, we present the first mountable ultrasonic microphones for use in studying mouse ultrasonic vocalizations. Wearable or mountable microphones have aided work in birds [15] and bats [16], but adapting this technology for rodents has posed the challenge of finding an ultrasonic-capable microphone small enough to be comfortably carried by animals as small as laboratory mice. By identifying these lightweight microphones capable of recording USVs, and developing custom attachment hardware and circuitry, we were able to assign a high percentage (90%) of vocalizations to an individual mouse. Another group simultaneously developed these microphones for use in rats, by surgically implanting the microphone directly onto the nostril of the rat [18]. Notable differences compared to our setup include that theirs used a single miniature microphone for one animal in a pair instead of one for each animal, and that they relied on deep neural network models paired with manual correction to assign vocalization identity. Their study validates the use of these microphones in measuring rodent ultrasonic vocalizations, and our study provides a flexible, detachable microphone that is generalizable to multiple species, including mice, and a computationally simple identity assignment.

The primary benefits of our system are that these mini-microphones are extremely low-cost (*$*0.60 each at the time of writing), are easy to implement – requiring very little material and computational investment – and are very customizable. We provide our designs of the circuit board and arm attachment, along with the materials list and procedures used. Utility of these mini-microphones also lies in their ease of integration with other systems. We demonstrated ready integration with video data. Our system has the potential to pair with an online credit-assignment system, behavioral perturbations, electrophysiology, or optogenetics, which would allow in-depth investigation of multi-animal vocal behavior. While our method allows easy and high-fidelity credit-assignment, existing array-based methods still have the benefit of featuring fully wireless behavior, which can accommodate animal recordings beyond a pair with greater ease. However, future versions of our system could involve a wireless setup, with appropriate high-speed data streaming or data logging.

Our findings are consistent with some found in array microphone systems [10,11,12], namely that female-assigned vocalizations often occur at very close range. As the largest proportion of unassigned USVs is in the similar close range, it is possible that some of them were produced by the females. Our finding that the majority of male USVs occur during ano-genital proximity (presumably with sniffing and stimulation) is consistent with the idea that the male’s UVSs directed to a female is a courtship-facilitation behavior.

We were able to perform identity assignment for a high percentage of vocalizations with only two microphones and very simple computations. However, further innovations with this system might enable an even higher rate of USV assignment. These future directions might include placing a mini-microphone on both sides of an individual mouse’s head; providing a second equidistant microphone would likely aid in assigning vocalizations produced at close snout-to-snout proximity but at an angle. A non-wired or non-surgical harness would also allow more naturalistic conditions for the vocal and social behavior, and would permit even longer recording sessions without complications from cable tangling. The video recordings may not only be useful to investigate behavioral components that are coordinated with mouse vocal production behavior, but could also serve as an additional source of triangulation information. Finally, we intentionally kept the identity-separation computations as simple as possible. We encourage other groups to experiment with variations on the system that may further improve its credit assignment capabilities.

