## Supplementary Materials for "Mountable miniature microphones to identify and assign mouse ultrasonic vocalizations"

### Materials and Methods

**Mouse Lines**

Adult male and female C57BL/6J mice were purchased from Jackson Laboratories (JAX #000664), and were 3-5 months of age at the time of recording. Prior to the surgical procedure, mice were group-housed in 5 animals per cage of the same sex, then individually housed after the procedure and between recording periods. The mice had free access to food and water. All animal treatment adhered to standards laid out by National Institute of Health guidelines, with approval from The Rockefeller University Institutional Animal Care and Use Committee (IACUC).

**Mini-microphone assembly**

We applied solder paste to the pads of each printed circuit board (PBC) with a custom stencil (PCBWay). We placed a JST connector, 0.1µF capacitor, and mini-microphone (Knowles SPU0410LR5H-QB) on their corresponding pads and ran through a reflow oven to affix the parts. We confirmed appropriate electrical connectivity of each mini-microphone PCB with a multimeter. We then glued a custom 3D-printed arm to the back of the PCB (For 3D-print and circuit files, see Supplementary materials).

**Mini-Microphone Attachment Surgery**

To securely attach the custom mini-microphone, we used a locknut and screw design to allow convenient attachment and removal of the  mini-microphone as needed between recording sessions. A steel serrated flange locknut (Zinc-Plated, 4-40 Thread, 1/4" Wide from McMaster-Carr) was placed on the head of the mice in a minimally invasive surgical procedure. Aseptic techniques were observed throughout the surgery, and surgical tools and the locknut to be implanted were steam sterilized using a 34-minute autoclave gravity cycle at 121°C. For the surgery, anesthesia was induced using isoflurane and the mouse was placed in a stereotaxic frame, secured with a nose cone and ear bars. A heating pad was placed under the mouse body to maintain appropriate body temperature throughout the procedure, and an ophthalmic ointment was applied to the eyes. Hair on the top of the head was trimmed and removed using a small amount of hair removal cream. The exposed skin was cleaned with an iodine swab stick then a sterile alcohol prep pad, repeated three times. An incision was made on the skin above the skull, creating an opening running anterior-posterior over the skull, roughly between the frontonasal suture on the anterior portion and lambdoid suture on the posterior portion. A hemicycle cut was made along either side of the opening to create an oval skin opening centered approximately on bregma. The membranes on the skull were removed using a combination of scraping with fine surgical forceps and application of a small amount of 3% hydrogen peroxide. As an analgesic to help with postsurgical recovery, a small amount of 0.25% Bupivacaine (2.5mg/mL) was applied to the skull and edges of the skin opening using a syringe. Once the opening was cleaned with bacteriostatic 0.9% sodium chloride saline and dried, 3M Vetbond was used as a biocompatible adhesive to seal the skin opening to the skull.

After the Vetbond dried, the locknut was secured to the surface of the skull using a small amount of Loctite Superglue, centered on the oval skin opening. To further stabilize the locknut and cover all exposed skin, we used OrthoJet dental cement to cover the flange surrounding the locknut and all skin edges. Three layers of OrthoJet were applied, ensuring the next layer is applied only when the preceding one had dried completely. The mouse was then placed in a clean cage on a heating pad for recovery, provided with hydrogel and food, and closely monitored in the next days for potential complications. The procedure used was identical between male and female mice.

**Behavioral Paradigm**

Recording sessions were conducted with a male and female mouse together, after both had received the locknut implant described above and recovered appropriately. During a recording session, we connected 3 separate channels to the Ultrasound Gate 416H (USGH) recording interface: the male mini-microphone, the female mini-microphone, and an overhead microphone (Avisoft Bioacoustics CM24/CMPA) suspended on the top of the acrylic box, 15 inches from the floor of the box. Mini-microphones were connected to the USGH via a 5-pin XLR to screw terminal adapter (Sescom). We used the custom skull attachment to secure the custom flexible printed circuit board (PCB) (PCBWay) with the Knowles MEMS microphone (see Supplementary material for circuit design). The 15 inches from the bottom ensures that the overhead microphone captures the full spatial coverage of the base surface area (12x12 inches) of the box. All audio recordings were conducted using the Avisoft-RECORDER software (Avisoft Bioacoustics) on a Windows PC. Video data was recorded from below using the Basler Ace 2 camera (a2A1920-160ucPRO), and from above with a Firefly S camera (Teledyne FLIR). Bonsai was used to record and process videos. Video and audio recordings received simultaneous 1 Hz pulses to enable post-hoc synchronization.

Prior to the mouse social USV recording, the 2 mini-microphones were gain-matched to ensure a comparable level of sensitivity to the audio being received. The two mini-microphones were attached to a wall of the recording box next to each other, equidistant to an Ultrasonic Dynamic Speaker (Vifa #60108, Avisoft Bioacoustics) connected to a 216H Ultrasound Playback Interface. An audio file of sample ultrasonic vocalizations was played, while the Avisoft-RECORDER spectrograms for the two mini-microphones were visually monitored and the gain for each respective spectrogram channel was adjusted to a close match. A short recording from the mini-microphones was then recorded so it could be used as a reference point during data analysis.

To secure the PCB with the mini-microphone to the head mount on the mice, the mice were first briefly anesthetized using isoflurane. The PCB, glued to a custom 3D printed acrylic mount was placed onto the locknut on the mouse's head, and secured using a 4-40 Phillips head screw (McMaster Carr; See Supplementary Materials for 3D designs). The PCB and mini-microphone were orientated with the acoustic port facing directly towards the mouse's mouth (0° angle, see Fig 2b). The mouse was then moved to the recording setup and the mini-microphone wire was attached to the sound channel; the mouse was given around 10 minutes to recover from anesthesia and acclimatize to the environment. The procedure was identical for both male and female mice. The audio and video recordings were started once the female's microphone wire was connected and she was placed inside the recording box, after having connected the male mouse. A 40-minute recording was conducted in this social context, with constant monitoring to ensure the cables were not tangled as to impede the free movement of either mouse in the box. If cables began to tangle, we manually rotated a small 3D printed piece (Fig 3a) through which both mini-microphone cables pass in separate channels; this rotation allowed any tangling to transfer elsewhere on the wires. So far this has been a workable solution for pairs of 2 mice even over an hour of recording. Once the recording session was complete, the mice were unplugged from their cables, and the screw loosened to remove the acrylic mount / PCB attachment. The mice were then returned to their individual housing cages until the next session.

**Audio/Video alignment**

An AdaFruit Feather M0 Basic was programmed to generate 1Hz digital pulses, with a length randomly selected from 100ms, 200ms, 300ms, 400ms, and 500ms. For videos, the digital pulses were delivered to two LEDs that were positioned within clear view of the top and bottom camera, respectively. For audio, the digital pulses were delivered to the digital input port on the Avisoft USGH 4-channel recorder used to collect recording from all microphones. For each recording modality, frames containing pulse starts and ends were identified. We confirmed the pulse patterns matched, then created a global time frame that progressed by 1 second with each pulse start. For each 1-second block, for each modality, the number of F frames was counted, and individual frames were assigned a time increment 1/F. Thus audio and video were both aligned to the same uniform external source and every frame was placed within a shared global time frame.

**Video tracking and inter-animal distance measurements**

We trained a multi-animal bottom-up SLEAP model (Pereira et al. 2022) on 527 total frames across seven videos taken from the bottom view camera (see Fig 3a, 4a). We labeled 11 parts (nose, neck, left ear, right ear, body center, tail base, middle tail, right forepaw, left forepaw, right rear paw, left rear paw) but for this study we only used the nose, neck, body center, and tail base. After training was completed, we ran model inference for each video, and manually corrected any identity swaps between male and female skeletons. We measured the left-to-right and front-to-back lengths inside of the recording arena in centimeters, and in pixels with the first frame of each video. We used these pixel/centimeter ratios to convert animal distances output from SLEAP in pixels to centimeters.

**Analysis of identity assignment**

For each recording session, we generated spectrograms (fs=250e3, Hann window, nperseg=256) from the wav files for each channel (top, male mini, female mini). For all recordings, we used the spectrogram output and applied no additional filtering. We used the top recordings as a ground truth for acoustic and temporal properties, and the mini-microphone recordings to compare volume differences for each individual syllable to use for identity assignment. We used MSA 2.0 [17] (Stoumpou et al. 2023) to perform general acoustic and temporal analysis of the top microphone recordings for both the gain-matching session and courtship session. From MSA 2.0, we obtained the start and end times of each USV syllable on the top recording. Within each syllable, for each time bin, we identified the three frequencies that had the highest amplitude in the top microphone recorded spectrogram (Fig 3b). For that same time bin, we measured the mean amplitude across those three frequencies, in the male and female mini-microphone spectrograms. We calculated the median amplitude across all time bins for a single syllable, for male ($M_{amp}$) and female ($F_{amp}$) spectrograms.

Once we obtained male and female amplitudes for every USV, we took the log of the ratio between male and female: $\log\left({M_{amp}}/{F_{amp}}\right)$ for each syllable. Here, an amplitude ratio greater than 0 means that a single USV was louder on the male mini-microphone, while a amplitude ratio below 0 indicates that the USV was louder on the female mini-microphone. A amplitude ratio value of 0 signifies no volume differences.

The $\log\left({M_{amp}}/{F_{amp}}\right)$ amplitude ratio was computed, as above, for every syllable from the gain-matching session. We computed the mean ($gain_{mean}$) and standard deviation ($gain_{SD}$) across all syllables for a single gain-matching session. These values were used as a ground truth measurement of inherent amplitude differences between the two mini-microphones for each session. Because every courtship recording session was preceded by a gain-matching session, we used the same-day session as the amplitude ratio benchmark. We classified a single courtship $USV_x$, that had amplitude-ratio $\log\left({M_{amp}}/{F_{amp}}\right)_x$ according to the following logic:

| | |
|---|---|
| Male USV | $\log\left({M_{amp}}/{F_{amp}}\right)_x \geq gain_{mean} + gain_{SD}$ |
| Female USV | $\log\left({M_{amp}}/{F_{amp}}\right)_x \leq gain_{mean} - gain_{SD}$ |
| Unassigned USV | $gain_{mean} - gain_{SD} \leq \log\left({M_{amp}}/{F_{amp}}\right)_x \leq gain_{mean} + gain_{SD}$ |

### Supplementary Figures

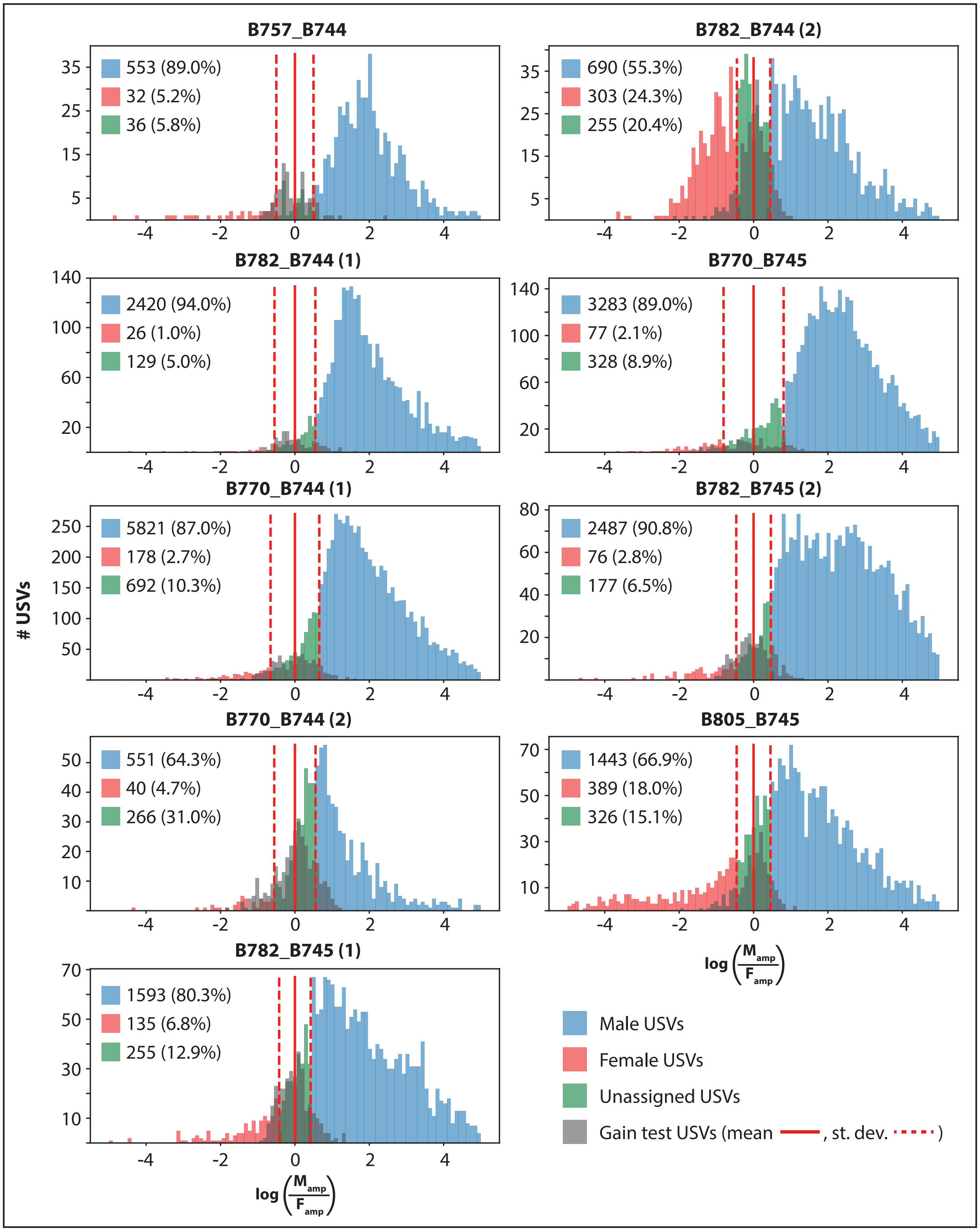

**Fig S1. Amplitude ratios across all sessions.**

Histograms displaying the male-female amplitude ratio of all syllables across all 9 40-minute recording sessions. Histograms are shown in the same format as Fig 3d. The distribution of male:female amplitude ratios across the gain-matching session (gray, solid red line: mean; ± 1 standard deviation: dashed red line) is used as the point of comparison for amplitude differences in the courtship session. Male-assigned (blue), female-assigned (red), and unassigned syllables (green) based on difference from gain-matching session mean.

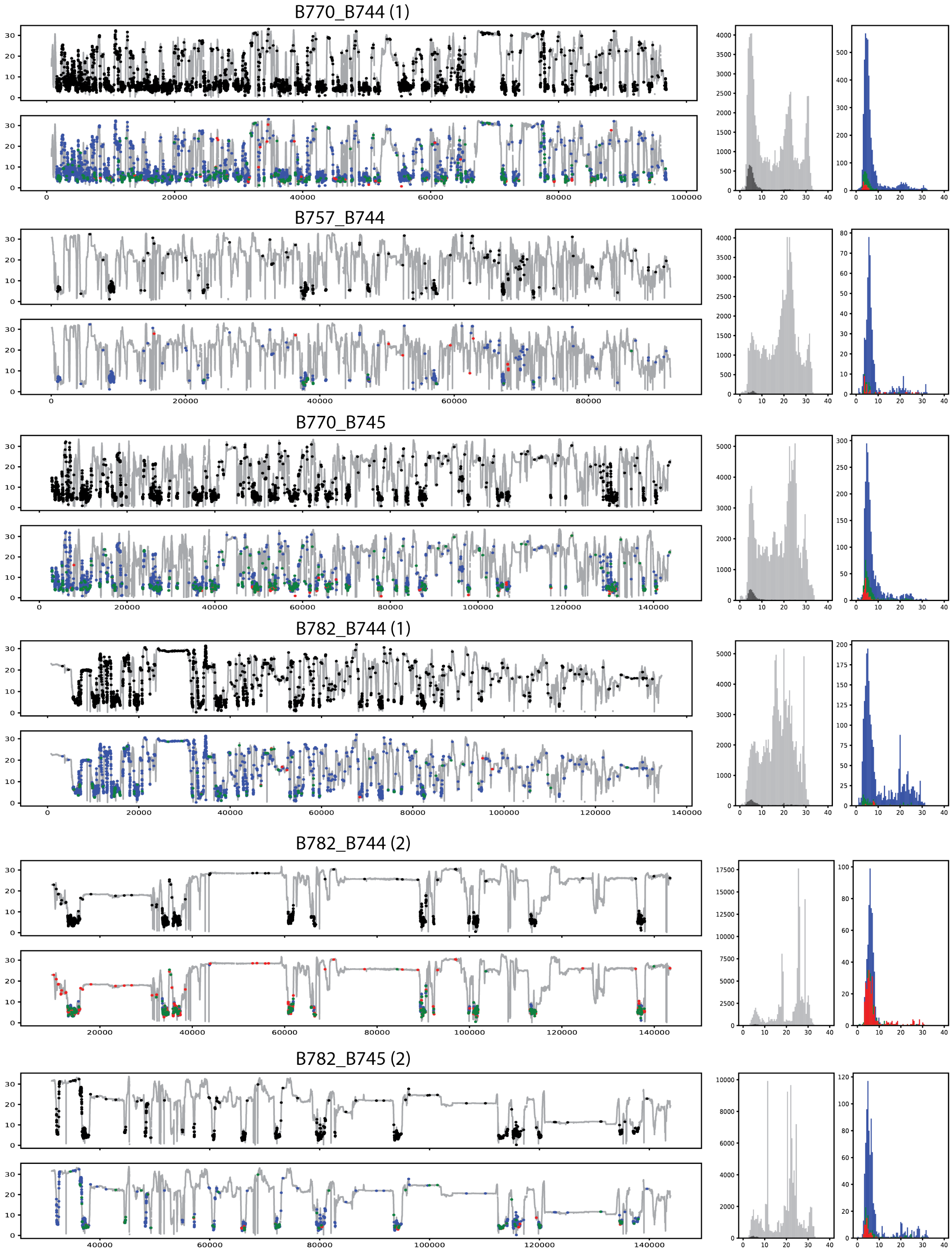
B770_B744 (1)
B757_B744
B770_B745
B782_B744 (1)
B782_B744 (2)
B782_B745 (2)

**Fig S2. Comparing inter-animal distances during USV production.**

Plots show distribution of neck $_M$ – neck $_F$ distance during frames containing, from left to right, male vocalizations, female vocalizations, unassigned vocalizations, all vocalizations, randomly-selected non-vocalizing periods, and all non-vocalizing periods. Data shows n = 6 individual sessions, with the aggregate data presented in main text Fig 4c.

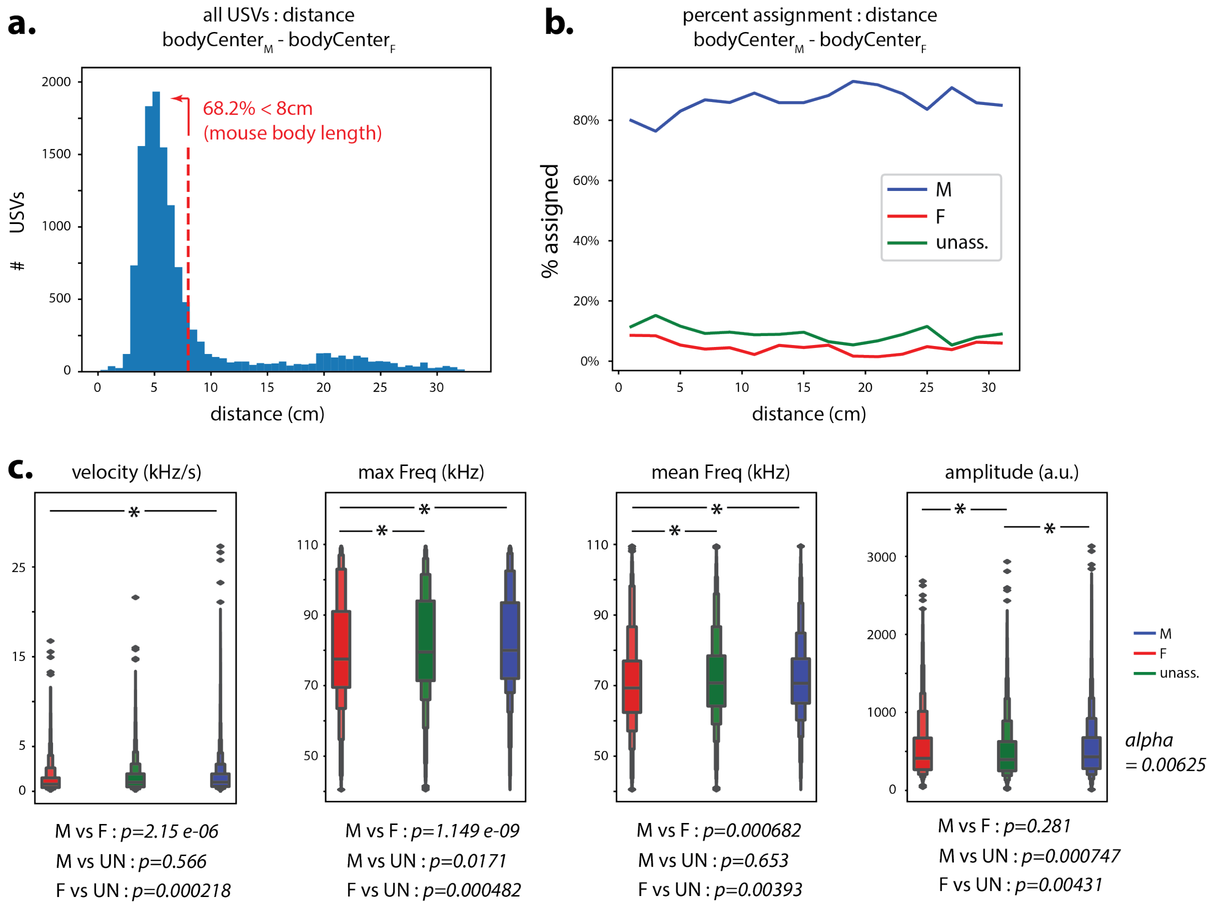


**Fig S3: USV distance and acoustic parameters.**

(a) Histogram of number of USVs that occur at different inter-animal distances between $bodyCenter_M$ and $bodyCenter_F$. Red line: 8cm (approx. adult mouse length). (b) Proportion of USVs assigned to male (blue), female (red), or unassigned (green) at different inter-animal distances. (c) Acoustic properties of male (blue), female (red), and unassigned (green) vocalizations. Below, p-values from independent t-tests, Bonferroni corrected.

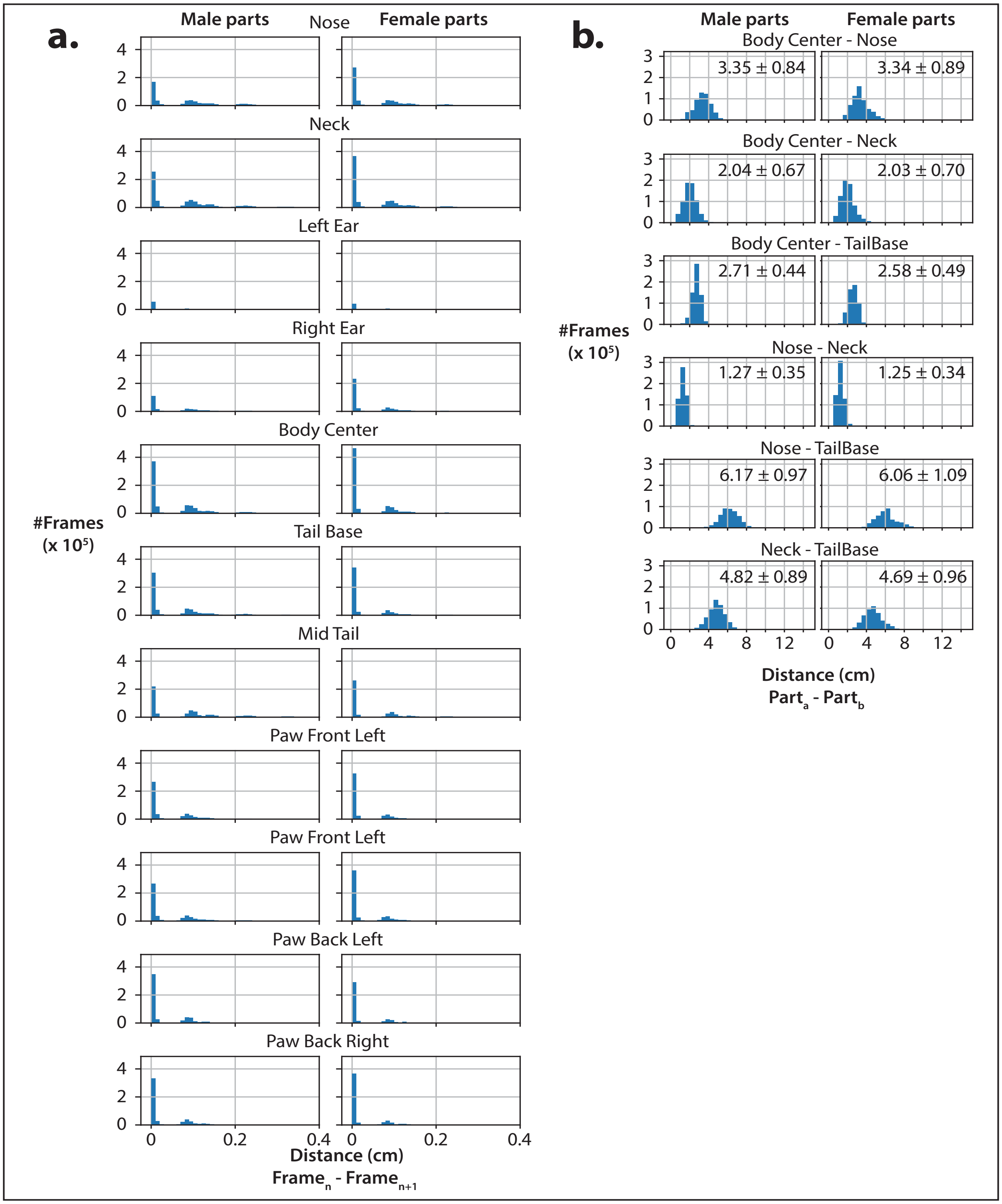

**Fig S4: SLEAP tracking validation.**

(a) Histogram of frame-to-frame distances (cm) for each individual body part. N=6 sessions. Bimodal distributions demonstrate both stationary epochs (peak close to 0 cm) and movement epochs (peak close to 0.1cm). All substantially less than the length of 1 adult mouse, ~8cm. (b) Histogram of part-to-part distances (cm) along main body axis. For each distribution, mean ± standard deviation in black text inset (All less than the length of 1 adult mouse, ~8cm). N=6 sessions.

### Acknowledgements

We thank the Rockefeller University Precision Instrumentation & Technology facility and Dr. Jazz Weisman for help with circuit board design and development, and the Rockefeller University Comparative Biosciences Center for excellent care of our animals. Peter Schade, Dr. César Vargas, and Hector Bermudez-Rivera provided valuable feedback and discussion.

### Author Roles

ENW and EDJ conceived of the experiment. ENW and VHYY developed custom hardware. ENW, VHYY, and WCD performed surgeries, collected behavioral data, and performed data analysis. ENW, VHYY, and EDJ wrote the manuscript.
